# Boron Supply Restores Aluminum-Blocked Auxin Transport by Modulation PIN2 Trafficking in the Arabidopsis Roots

**DOI:** 10.1101/2021.12.09.471968

**Authors:** Lin Tao, Yingming Feng, Yalin Li, Xuewen Li, Xiaodong Meng, Meng Li, Niloufar Pirayesh, Sakil Mahmud, Sergey Shabala, František Baluška, Lei Shi, Min Yu

## Abstract

This study tested a hypothesis that boron (B) supply alleviates aluminum (Al) toxicity by modifying auxin distribution in functionally different root zones. Auxin distribution and transport at various Al and B ratios were analyzed using the range of molecular and imaging techniques. Al stress resulted in increased auxin accumulation in root apical meristem (MZ) and transition zones (TZ) while reducing its content in elongation zone (EZ). This phenomenon was explained by reduction in basipetal auxin transport caused by Al blockage of PIN2 endocytosis, regulated at posttranscriptional level. This inhibition of PIN2 endocytosis was dependent on actin filaments and microtubules. B supply facilitated the endocytosis and exocytosis of PIN2 carriers via recycling endosomes conjugated with IAA to modify Al-induced auxin depletion in the EZ. However, disruption of auxin signaling with auxinole did not alleviate Al-induced inhibition of root growth. B supply alleviates Al-induced inhibition of root growth via restoring the endocytic recycling of PIN2 proteins involved in the basipetal (shootward) auxin transport, restoring Al-induced auxin depletion in the elongation zone.

**Short summary:** Aluminum-intensified PIN2 abundance, nontranscriptional, via repressing PIN2 endocytosis to block polar auxin transport, and this adverse effect could be alleviated by boron supply.

## Introduction

Aluminum (Al) toxicity is a major limiting factor for plant growth and development in acidic soils throughout the world (von Uexküll and Mutert, 1995), which executes potent inhibitory effect on root growth generally within less than one hour of Al exposure (Kopittke et al., 2015). Numerous studies have reported that the root apical transition zone (TZ), located between the apical meristem (MZ) and basal elongation zone (EZ) (Verbelen et al., 2006; Baluška et al., 2010; Baluška and Mancuso, 2013), is regarded as the Al-sensitive site in several species such as Arabidopsis thaliana (Illéš et al., 2006; Yang et al., 2014), maize (Eticha et al., 2005), rice (Wu et al., 2014) and sorghum (Sivaguru et al., 2013). Moreover, it has been suggested that the high sensitivity for the TZ in maize root apices to Al stress is linked to the basipetal auxin transport and responses (Kollmeier et al., 2000). Several studies have reported various Al-induced abnormal physiological processes, such as depolymerization of F-actin (Shen et al., 2008), elevated reactive oxygen species (ROS) levels (Sivaguru et al., 2013) and disturbance to endogenous ethylene and auxin hormonal profiles in root apices (Sun et al., 2010; Yang et al., 2014). However, the underlying mechanisms of the Al-induced inhibition of root growth remain to be elucidated.

Auxin (indole-3-acetic acid, IAA), a major plant phytohormone, plays a critical role in modulating plant growth and development (Friml, 2003; Zhao, 2010). Previous studies have reported that that Al inhibits the polar auxin transport (PAT) in the root apices, ultimately leading to auxin accumulation in the root cap, the root apical MZ and TZ (Kollmeier et al., 2000; Sun et al., 2010; Yang et al., 2014). Local application of IAA in the EZ of maize root tips ameliorated the inhibitory effect of Al toxicity on root growth (Kollmeier et al., 2000), implying that Al exposure might block basipetal auxin transport from the TZ to EZ in root tips in response to Al toxicity. However, the mechanistic basis of the Al-induced disruption of basipetal auxin transport and auxin distribution in root tips remain unknown.

Auxin transport in plants is mediated by auxin-influx carriers from AUXIN1/LIKE AUX1 (AUX/LAX) (Péret et al., 2012; Yang et al., 2006) and PINFORMED (PIN) families (Adamowski and Friml, 2015), along with the members of the ATP-binding cassette subfamily B (ABCB) for directing auxin transport (Geisler and Murphy, 2006). Moreover, the continuous cycling of PINs that occurs between the plasma membrane and endosomal compartments has been suggested to play a critical role in a polar auxin transport in root tips (Shibasaki et al., 2009; Schlicht et al., 2006). Currently, the most accepted model is that PIN carriers execute their functions on auxin-efflux at the plasma membranes. However, previous studies have reported that Al stress not only inhibited the endocytosis of PIN2 proteins (Shen et al., 2008), but also caused auxin accumulation within the epidermal and cortical cells of the Arabidopsis root apical transition zones (Sun et al., 2010; Yang et al., 2014), subsequently, leading to the arrest of root elongation. The latter phenomenon was attributed to Al-blocked basipetal auxin transport form the TZ to EZ in root apex, resulting in auxin accumulation in the former and depletion in the latter root apical zones (Kollmeier et al., 2000). However, no direct evidence was provided to support this hypothesis. Thus, the underlying mechanism whether Al-induced inhibition of root growth via modulating PIN2-dependent auxin transport requires further elucidation.

Acid soils are not only characterized by elevated Al^3+^ levels but also known to be deficient in boron (B) (Shorrocks and Bureau, 1997). Boron is an essential micronutrient for plant growth and development (Tanaka and Fujiwara, 2008; Camacho-Cristóbal et al., 2015) which is involved in plethora of physiological processes. One of them is in facilitating the formation of pectin gels via binding with rhamnogalacturonan II (RG-II) in the primary cell walls (O’Neill et al., 2004). Insufficient B levels results in major alterations in the cell wall and plasma membrane structure and function (O’Neill et al., 2004), the perturbation of auxin distribution in the root apex (Camacho-Cristóbal et al., 2015), repression the endocytosis of external cell-wall pectin (Yu et al., 2002), and slow root elongation (Yu et al., 2009; Li et al., 2018). Previous studies have reported that B deprivation-induced increase in the auxin content in stelar cells, as well as in the epidermal and cortical cells of the meristem zone of Arabidopsis (Camacho-Cristóbal et al., 2015). It has been suggested that B deprivation affects auxin transport carriers AUX1 and PIN2 to arrest of root elongation (Camacho-Cristóbal et al., 2015).

Interestingly, both Al toxicity and B deprivation evoke similar physiological processes such as root growth inhibition and disruption of auxin distributions in root apices (Yang et al., 2014; Sun et al., 2010; Camacho-Cristóbal et al., 2015). Moreover, numerous studies have reported that B supply alleviates the inhibitory effects of Al toxicity on plant growth and development. Two underlying mechanisms beyond this phenomenon were alteration of cell wall pectin contents and characteristics (Yu et al., 2009) and elevation of apoplast pH (Li et al., 2018). However, to the best of our knowledge, ameliorating effects of B on Al toxicity have not been attributed directly to auxin transport and signaling.

Here, we show that Al-induced upregulation of *PIN1* expression level may promote the shoot-derived auxin supply and enhance PIN2 membrane signals by inhibition of PIN2 intracellular trafficking. In the presence of Al, PIN2 has longer residence times at the plasma membrane, ultimately leading to auxin accumulation in the MZ and TZ root apex zones, while lower auxin levels were scored in cells of the EZ. Importantly, B supply not only downregulated PIN1 carrier abundance, but also efficiently facilitated the endocytosis and exocytosis of PIN2 through recycling endosomes to promote PIN2 endosomes-dependent auxin transport for restoration of the Al-induced shortage of auxin amounts in the root apical EZ.

## Results

### B Supply Ameliorates Al-Induced Inhibition of Root Growth and Reduces Al Accumulation in *Arabidopsis* Roots Apices

Previous studies have revealed that B supply predominantly alleviated Al-induced inhibition of root growth in pea (*Pisum sativum*) (Yu et al., 2009; Li et al., 2018). In this study, we also observed that B deprivation or Al toxicity could arrest of *Arabidopsis* wild type (*col-0*) root growth, and this adverse effect was attenuated by B supply (Figure 1A and 1B). Subsequently, we analyzed Al contents and distribution by using the morin dye (specific for active Al). The transition zone (TZ) of wild type (*col-0*) root apices had the highest Al-morin fluorescent signals (Figure 1C and 1D) and showed a strong time-dependency on Al exposure (Figure 1D). The presence of B efficiently reduced Al accumulation in the TZ of wild type root tips and mitigated Al toxicity.

**Figure 1.**
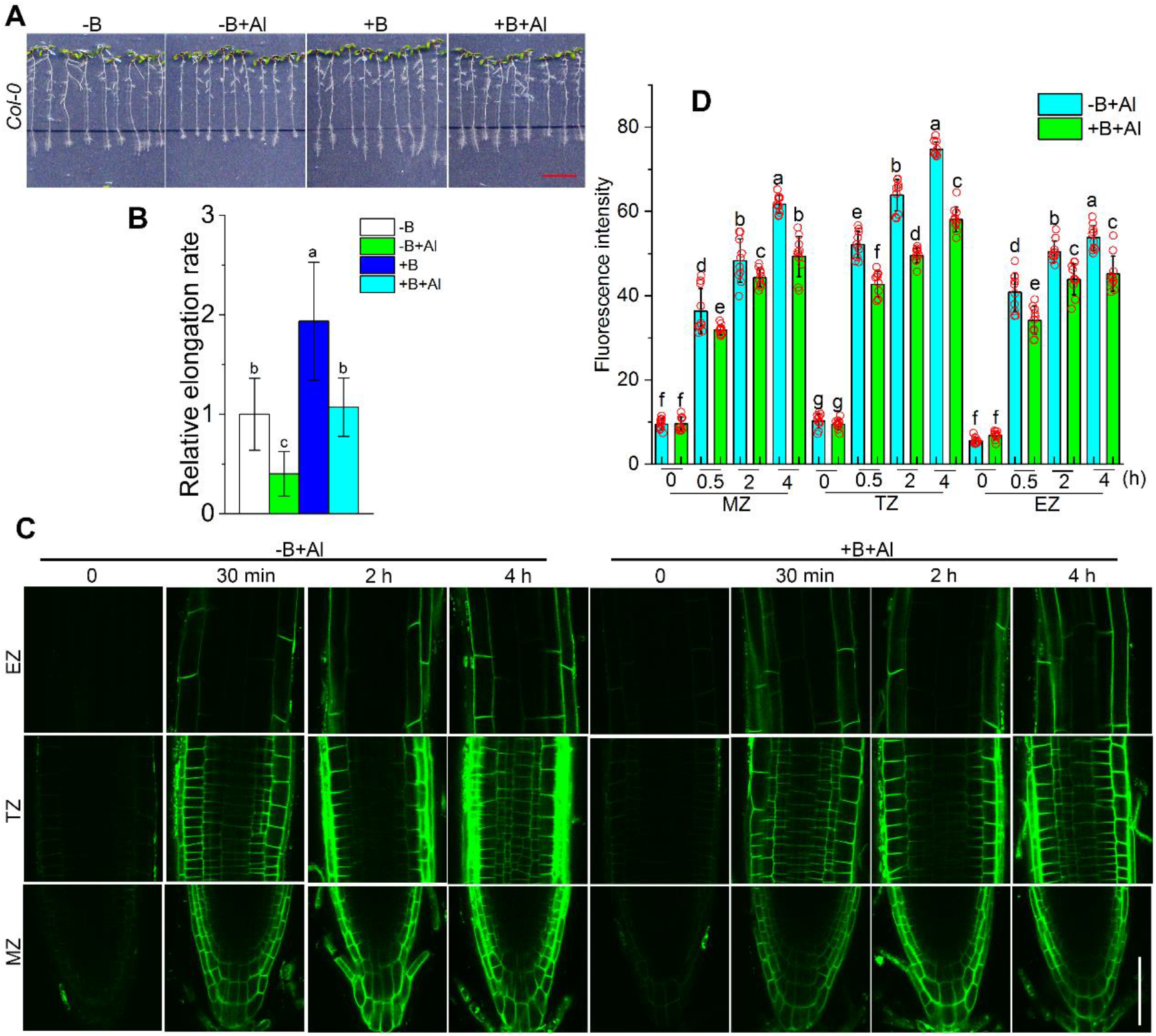
Effects of B and Al on *Arabidopsis* root growth and Al distribution. Six-day-old *Arabidopsis* (*col-0*) seedlings were treated with newly agar containing Al (0 or 100 μM AlCl_3_) and B [0 or 50 μM B(OH)_3_)] at pH 4.5 for 2 d. Subsequently, treated seedlings (> 20) were imaged and representative photos were shown in **(A)**. The dashed black lines indicate the initial location of root tips on new culture dishes. Relative elongation rate **(B)** was calculated and -B treatment was set to 1. Values are means ± SD (*n* = 10). **(C)** Six-day-old *Arabidopsis* (*col-0*) seedlings were further treated with solution containing Al (0 or 100 μM AlCl_3_) and B [0 or 50 μM B(OH)_3_)] at pH 4.5 for 0, 30 min, 2 h and 4 h, respectively. Subsequently, treated seedlings (> 20) were stained by 0.01 % morin dye for 5 min and then imaged by confocal laser scanning microscope (with one of 8-10 representative images shown in the panel). **(D)** the fluorescence intensity of Al-morin signal shown in **(C)**. Values are means ± SD (*n* = 10). Red circles represent individual data. Data labelled with different letters are significantly different at *P* < 0.05 (LSD’s test). Bars are 1 cm for **(A)** and 100 μm for **(C)**, respectively.

### B Supply Restores Al-Induced Decrease of Auxin Content in the Elongation Zone

The dynamic changes of auxin distribution patterns in cells were mapped using the auxin *DII-VENUS* sensor (where auxin content correlates negatively with the intensity of *DII-VENUS* signals) (Brunoud et al., 2012). In the absence of Al, the *DII-VENUS* fluorescent signal gradually declined from the MZ to EZ in primary root epidermis, regardless of B supply (Figure 2A). The B-deprived roots had a lower *DII-VENUS* fluorescent signal than that with adequate B supply in the MZ and TZ while no significantly difference was observed in the EZ (Figure 2B). This suggests that B deprivation leads to a higher auxin accumulation in the root apices. In the presence of Al, B-deprived roots had a weak *DII-VENUS* fluorescent signal in the MZ and TZ along with a strong signal in the EZ, while B-supplied roots had a weak *DII-VENUS* fluorescent signal in the whole root tips (Figure 1A), implying that Al-induced decrease in the auxin content in the EZ could be restored by B supply.

**Figure 2.**
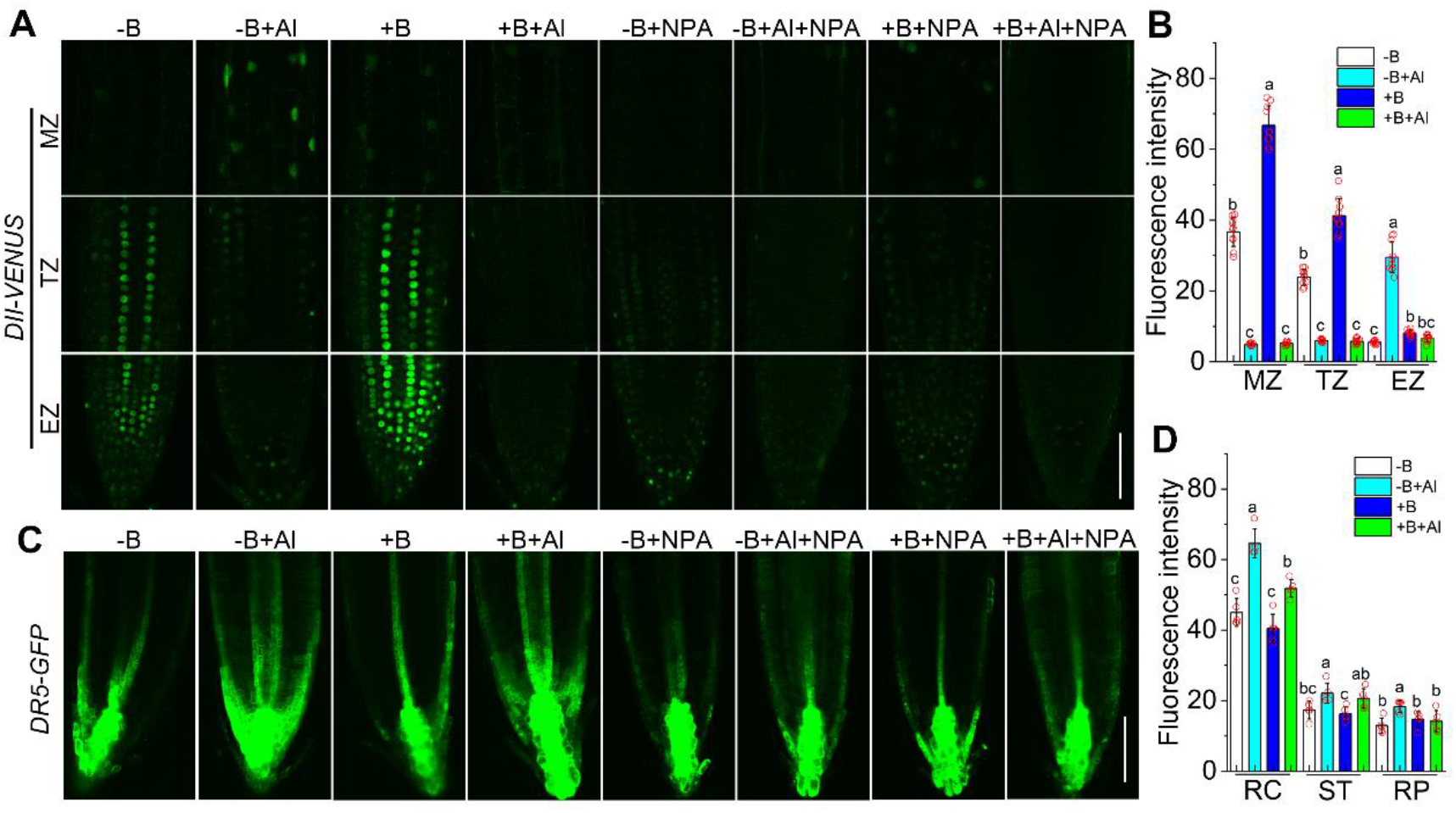
Effects of Al and B on auxin distribution patterns in *Arabidopsis* roots. Six-day-old transgenic *DII-VENUS* and *DR5-GFP* seedlings were treated with solution containing Al (0 or 100 μM AlCl_3_), B [0 or 50 μM B(OH)_3_)] and NPA (0 or 10 μM) at pH 4.5 for 4 h. The primary roots of transgenic *DII-VENUS* **(A)** and *DR5-GFP* **(C)** seedlings were then analyzed by confocal laser scanning microscopy (CLSM). **(B)** and **(D)** are the fluorescence intensity of **(A)** and **(C)**, respectively; values are means ± SD, *n* = 10 for **(B)** or 8 for **(D)**. Three independent experiments were conducted, with at least 10 plants analyzed in each treatment. Red circles represent individual data. Data labelled with different letters are significantly different at *P* < 0.05 (LSD’s test). Bars are 100 μm in **(A)** and **(C)**, respectively. Abbreviations: MZ, meristem zone; TZ, transition zone; EZ, elongation zone; RC, root cap; ST, stele tissue; RP, root profile.

To account for the possible drawbacks of the spatial distribution patterns of *DII-VENUS* fluorescent signals in the root apices, we employed another transgenic *DR5-GFP* maker line. The auxin response sensor of *DR5* is mainly expressed in the quiescent center (QC), columella and vascular cells under normal conditions (Sabatini et al., 1999). In the absence of Al, *DR5-GFP* fluorescent signals mainly originated from the quiescent center (QC), columella, vascular and the epidermal cells of root apices (Figure 2C). Application of Al in the absence of B elevated *DR5-GFP* fluorescent signals in the MZ and TZ, while B supply reduced *DR5-GFP* fluorescent signal from the root cap and epidermis (Figure 2C and 2D). In the presence of Al, B-deprived plants had a higher *DR5-GFP* fluorescent signals compared with B-sufficient roots (Figure 2D). Both the Al-induced increase of *DR5-GFP* and *DII-VENUS* fluorescent signals were strongly diminished in the presence of NPA (an inhibitor of auxin efflux from a cell) (Figure 2A and 2C). Collectively, these results demonstrated that Al interferes with the polar transport of auxin, leading to auxin accumulation in the MZ and TZ, but its depletion in the EZ. B supply efficiently prevented Al-induced depletion in auxin content in the EZ.

### Differential Effects of B and Al on Auxin Transport Carriers in Root Apices

Transgenic *GFP* for *PINs* plants were used to visualize the spatial distribution and response levels of auxin transport-related proteins. In the absence of Al, B-deprived plants had a higher PIN1-GFP signals compared with B-sufficient plants (Figure 3A and 3C). PIN1-GFP signals were higher in the MZ relative to the TZ among differential treatments. Al-induced increase in PIN1-GFP fluorescent intensity in the MZ was in agreement with the result of PIN1-GUS (Figure S3A). This implies that Al might upregulate *PIN1* expression levels; a result which was then verified by qRT-PCR analysis of *PIN1* gene in root tips (Figure S4). The MZ and TZ had a higher PIN2-GFP signals than the EZ (Figure 3B and 3D). In the absence of Al, no significant difference in PIN2-GFP signals from the plasma membranes in differ root zones was observed, regardless of B supply (Figure 3B). Al exposure increased PIN2-GFP signals from the plasma membranes in the TZ and MZ regardless of B supply (Figure 3D). Similar results were reported for Al-induced increase of signals from PIN2-GUS plants (Figure S3B). No significant changes in the *PIN2* transcript levels were found in response to Al (Figure S4), hinting that Al-induced increase in PIN2-GFP signals might occur at posttranscriptional level. In addition to PIN1 and PIN2 carriers, AUX1 carriers also play a pivotal role in auxin influxes. Al treatment increased AUX1-YFP signals regardless of B supply (Figure S2A and S2B); however, *AUX1* gene expression levels were unaffected among differential treatments (Figure S4) once again suggesting a possibility of posttranscriptional regulation. In contrast to PIN1-GFP and PIN2-GFP signals, there were no difference in PIN3-GFP, PIN4-GFP and PIN7-GFP signals after Al exposure regardless of B supply (Figure S1A and S1B); this was further verified by qRT-PCR analysis of *PIN3, PIN4* and *PIN7* expression (Figure S4).

**Figure 3.**
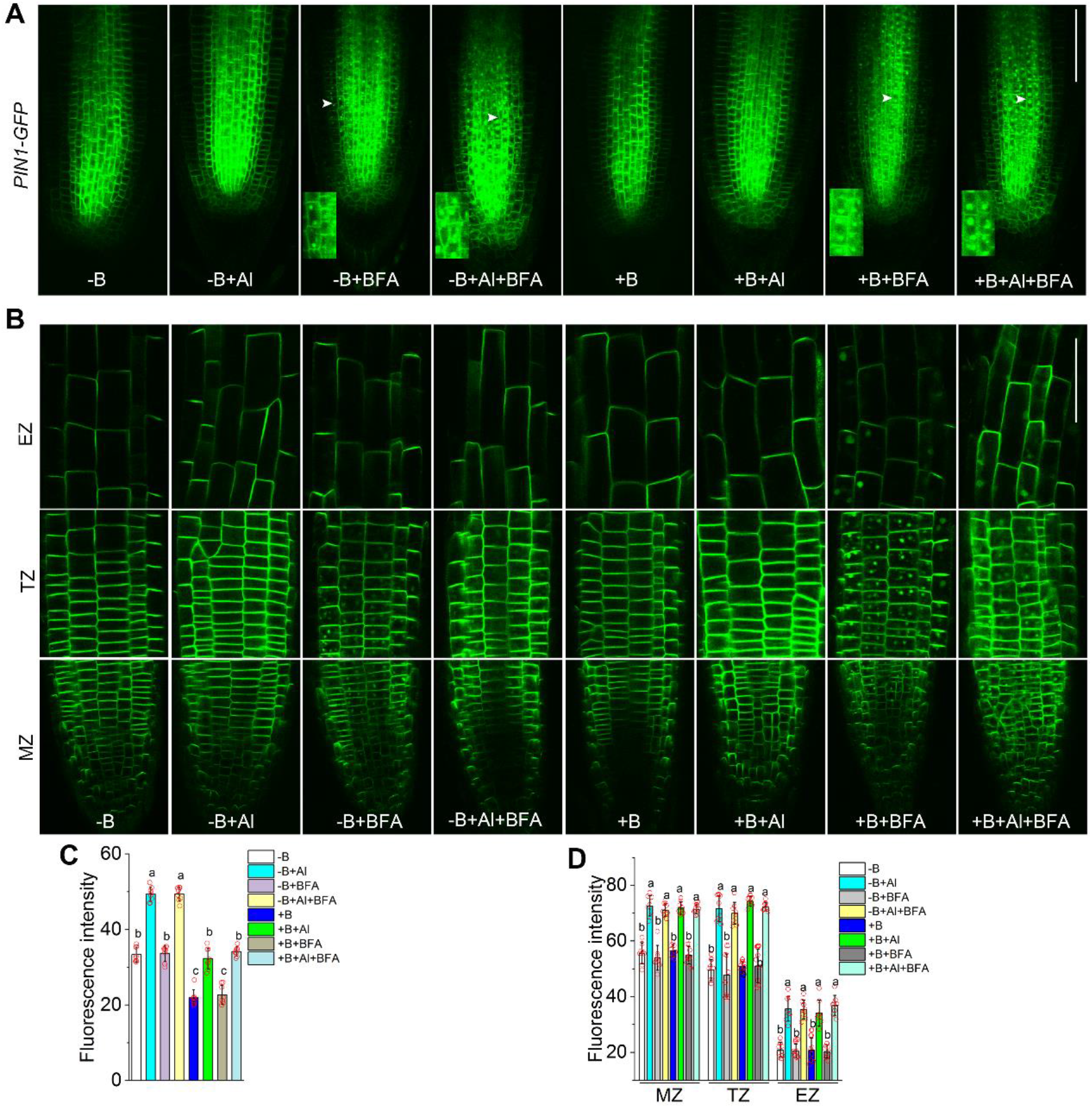
Effects of Al and B on the endocytosis of PIN1 and PIN2 carriers. Six-day-old transgenic *PIN1-GFP* **(A)** and *PIN2-GFP* **(B)** seedlings were treated solution containing Al (0 or 100 μM AlCl_3_), B [0 or 50 μM B(OH)_3_)] and BFA (0 or 100 μM) at pH 4.5 for 4 h. Treated seedlings (> 20) were imaged by CSLM. One (of 10) representative image is shown. **(C)** and **(D)** quantify the fluorescence intensity of PIN1 and PIN2 signals in **(A)** and **(B)**, respectively. Values are means ± SD (*n* = 8). Red circles represent individual data. Images in the lower corner of **(A)** are the enlarged locations indicated by white arrows. At least three independent experiments (with *n* > 10 individual plants in each). Data labelled with different letters are significantly different at *P* < 0.05 (LSD’s test). Bars are 100 μm in **(A)** and **(B)**, respectively.

Auxin transport carriers undergo continuous recycling between the plasma membrane and endosomal compartments. In this study, we employed BFA (brefeldin A, an inhibitor of exocytosis but not endocytosis) to decipher the endocytic trafficking of auxin transport-related carriers in root apices under varying conditions. In the absence of Al, and regardless of B supply, BFA-induced spots were found in PIN1, PIN2, PIN3, PIN7 and AUX1 lines (Figure 3A and 3B; see supplemental data Figure S1A and S2A). The BFA-induced spots were also detected in the upper cell layers of QC in transgenic *PIN4-GFP* plants (Figure S1A). Interestingly, the QC and columella cells displayed insensitivity to BFA reagents in transgenic *AUX1-YFP* (Figure S2A), *PIN3-GFP* (Figure S1A) and *PIN4-GFP* (Figure S1A) lines.

Noteworthy, it was clear that PIN2 endocytosis was severely repressed after Al exposure, while the other auxin carriers such as PIN1, PIN3, PIN7 and AUX1, were unaffected, implying Al selectively blocked the endocytosis of auxin transport-related carriers. Although B supply did not affect the expression pattern of PIN2 gene too (Figure S4), it has promoted the formation of BFA-induced PIN2 spots, especially in the TZ of root apices (Figure 3B). The number, size and signals intensity of BFA-induced PIN2 spots in the TZ of root tips was promoted by B supply. The EZ had the weakest fluorescent signals and lower frequency of BFA-induced PIN2 spots compared with TZ in all treatments, implying that PIN2 mainly operates in the root TZ.

Taken together, these data suggest that Al-induced increase in auxin content in root apices, especially in the MZ and TZ, may be causally related to upregulating PIN1 gene expression and repressing endocytosis of PIN2 carrier, leading to preferential auxin accumulation in MZ and reduced auxin transport into EZ. B supply efficiently promotes PIN2 endocytosis operating mainly in the root TZ thus mediating basipetal auxin transport via Al-sensitive domains in the TZ.

### Al-Induced Inhibition Endocytosis of Membrane PIN2 Carriers Depends on Posttranscriptional Regulation

To further distinguish between mechanisms of transcriptional or posttranscriptional regulation of PIN2 by Al, we employed cycloheximide (CHX, an inhibitor of protein synthesis) and cordycepin (COR, an inhibitor of transcription) used previously to decrease PIN1-GFP membrane signals (Geldner et al., 2001; Marhavý et al., 2011). Here, 50 μM CHX downregulated PIN2-GFP membrane signals in the TZ regardless of B supply (Figure 4A and 4B). In contrast, the simultaneously application of Al and CHX did not diminish PIN2 membrane signals. To rule out the possibility whether Al-induced intensification the re-secretion of PIN2 carriers to maintain the higher PIN2 membrane signals, we employed BFA reagent to repress the exocytosis process of PIN2 proteins in the presence of CHX. The results revealed that, in the absence of Al, BFA-induced PIN2 spots (Figure 3B) were abolished by CHX exposure regardless of B supply (Figure 4A). This implies that CHX disturbs the aggregation of BFA-induced PIN2 spots and discloses the facts of the direction and lytic degradation of PIN2 carriers to vacuoles. Conversely, the synergistic effect of CHX and BFA did not influence PIN2 membrane signals under Al stress indicating that Al-induced increase of PIN2 membrane signals was not affected by the exocytosis of newly synthesized PIN2 proteins, but by endocytosis of PIN2. Subsequently, we detected PIN2 membrane signals in the presence of 100 μM COR. COR exposure diminished PIN2 membrane signals that were also repressed after Al exposure regardless of B supply (Figure 4A and 4B). Overall, those findings suggest Al-induced increase of PIN2 membrane signals depends on posttranscriptional regulation.

**Figure 4.**
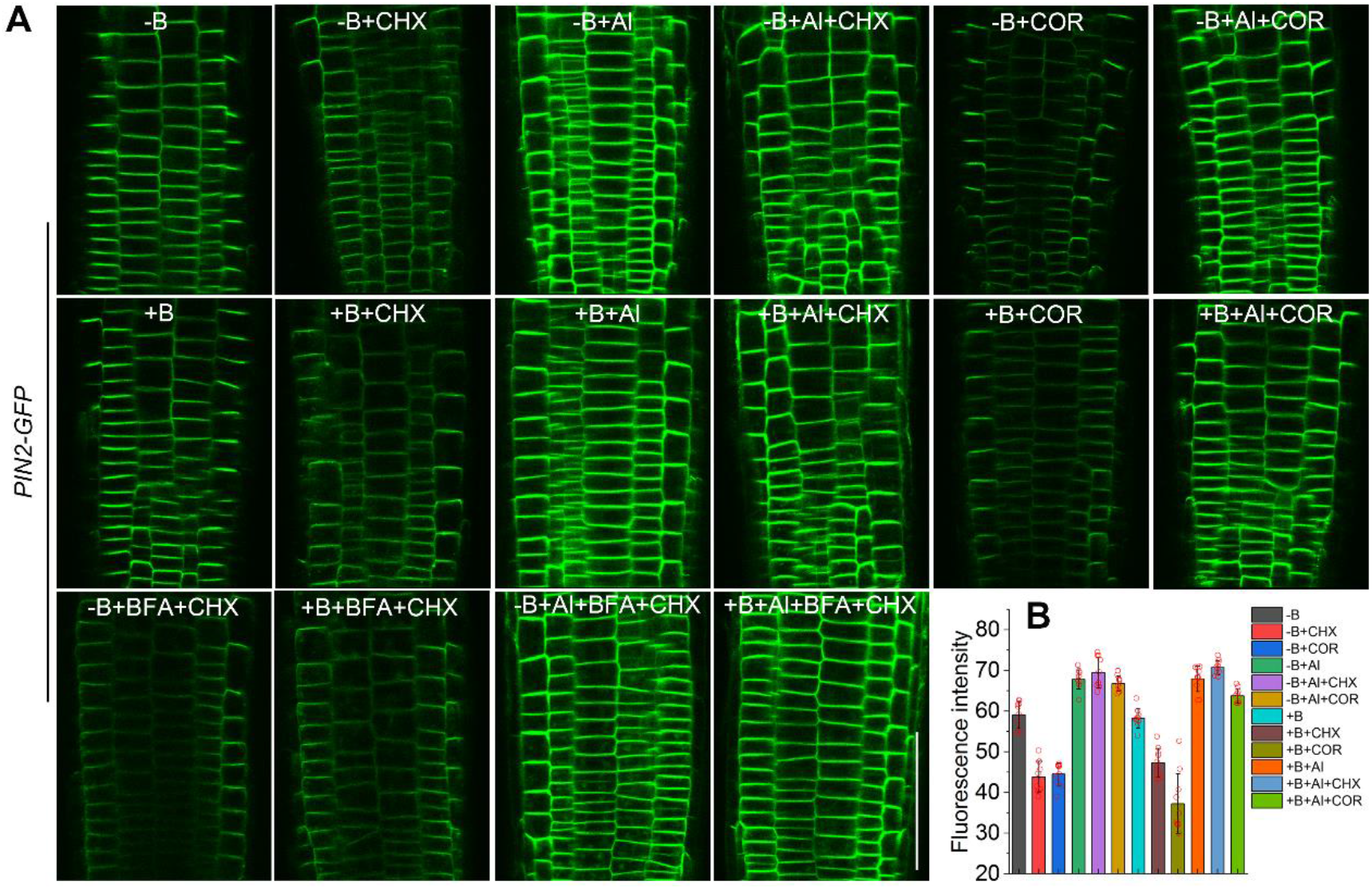
Al-induced increase of membrane PIN2 signals independently transcription and PIN2 proteins biosynthesis. **(A)** Six-day-old transgenic *PIN2-GFP* seedlings (> 10) were treated with solution containing Al (0 or 100 μM AlCl_3_), B [0 or 50 μM B(OH)_3_)], CHX (0 or 50 μM cycloheximide), COR (0 or 100 μM cordycepin) and BFA (0 or 100 μM) at pH 4.5 for 4 h. A typical CSLM images were then taken from transition root zones and **(B)** the fluorescence intensity of PIN2-GFP signals quantified. Values are means ± SD (*n* = 10). Red circles represent individual data. Bars is 100 μm.

### Al-Induced Inhibition of PIN2 Endocytosis Involves Actin Filaments and Microtubules

Previous studies have reported that stabilizing actin can interfere with endocytosis (Dhonukshe et al., 2008; Shen et al., 2008). Here, we employed actin-binding domain 2 of fimbrin gene fused with the green fluorescent protein gene (*ABD2-GFP*) line to analyze the effects of B and Al on the actin cytoskeleton. In contrast to Al-untreated roots, Al treatment led to the formation of thick actin bundles, which is a characteristic feature of actin stabilization (Dhonukshe et al., 2008), and B supply could ameliorate actin filament signals in plasma membranes and cytoplasmic spaces (Figure 5A and 5B). These observations indicate that B-supply may alleviate Al-caused stabilization of actin filaments.

**Figure 5.**
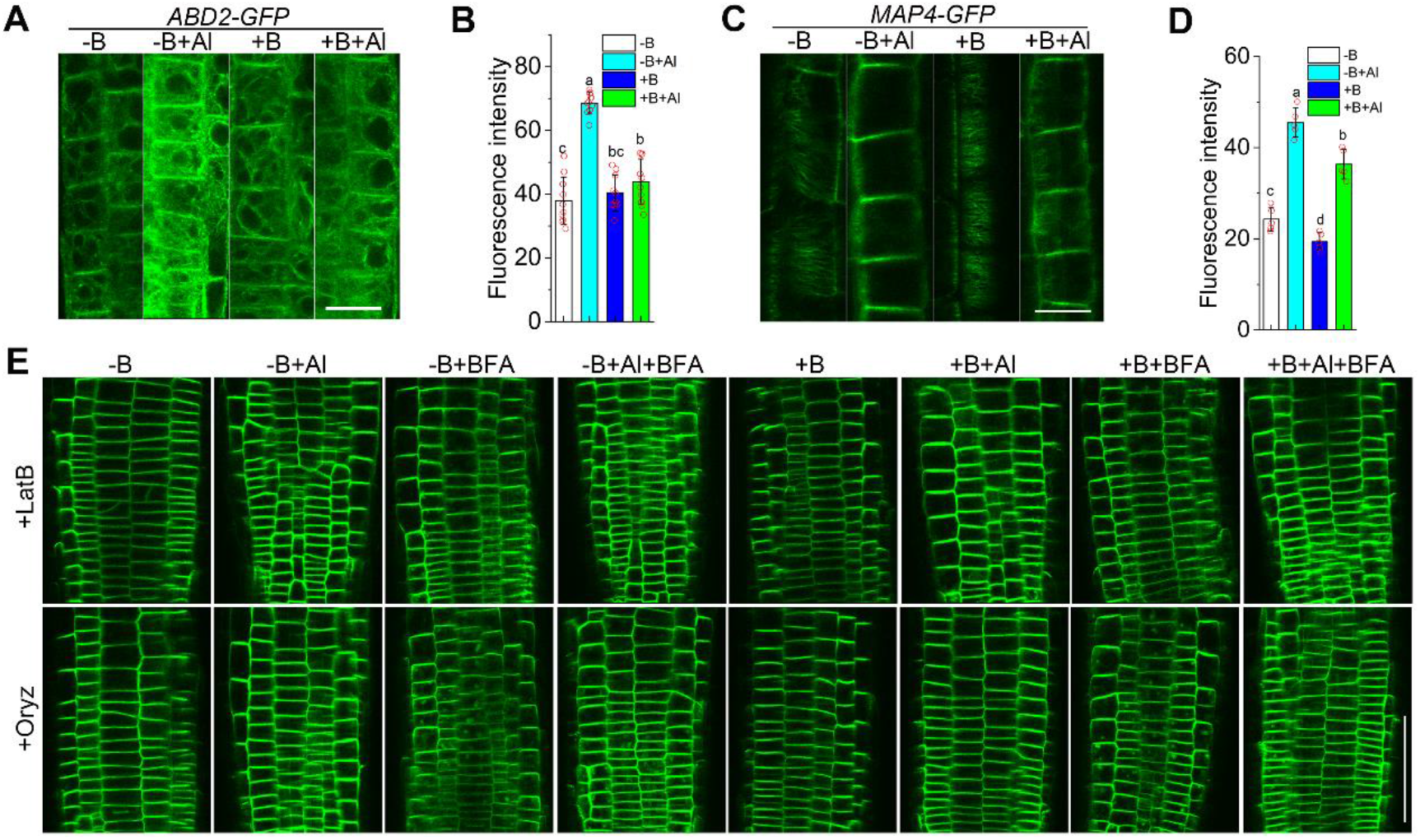
Effects of B and Al on actin filaments and microtubules in *Arabidopsis* roots. Six-day-old transgenic *ABD2-GFP* **(A)** and *MAP4-GFP* **(C)** seedlings were treated with solution containing Al (0 or 100 μM AlCl_3_) and B [0 or 50 μM B(OH)_3_)] at pH 4.5 for 4 h. A typical CSLM images were then taken from transition root zones. One (of > 20) typical image is shown. **(B)** and **(D)** the average intensity of the fluorescence membrane-GFP signals in shown in **(A)** and **(C)**, respectively. **(E)** Typical CSLM images of transgenic *PIN2-GFP* seedlings treated with 0.2 mM CaCl_2_ solution containing 10 μM LatB or 20 μM Oryz, Al (0 or 100 μM AlCl_3_) and B [0 or 50 μM B(OH)_3_)] at pH 4.5 for 4 h (transition zones). Bars are 20 μm for **(A)** and **(C)** and 100 μm for **(E)**, respectively. Data labelled with different letters are significantly different at P < 0.05 (LSD’s test).

Moreover, the movement of endocytic vesicles in the cell periphery depends on the actin filaments, whereas the fast movement of it in the central regions is mostly microtubule-dependent (Dhonukshe et al., 2008; Gasman et al., 2003). Therefore, we also employed microtubule-associated protein 4 gene fused with green fluorescent protein gene (*MAP4-GFP*) line to analyze the effects of B and Al on the microtubule cytoskeleton. In the absence of Al, microtubules were packed orderly in the cytosol regardless of B supply (Figure 5C). Similar to Al-caused changes for actin filaments (Figure 5A), Al exposure not only induced the formation of thick microtubules bundles in plasma membranes, but also weakened cytosol signals of microtubules (Figure 5C and 5D), implying Al treatment stabilized microtubules too. Boron deprivation aggravated this stabilization induced by Al toxicity (Figure 5C and 5D).

As expected, loss of actin filaments induced by latrunculin B (LatB) completely abolished the formation of BFA-induced PIN2 spots in root apical TZ (Figure 5E), indicating the role of actin filaments in the endocytosis of PIN2 carriers. However, the depolymerization of microtubules by oryzalin (Oryz) did not affect BFA-induced PIN2 spots formation, but weakened the size and fluorescent intensity of BFA-induced PIN2 spots (Figure 5E); moreover, compared with results in Figure (3B and 4A), the extent of Al-intensified PIN2 membrane signals was diminished (Figure 5E). These results suggest that Al-induced block of PIN2 endocytosis at the plasma membranes depends not only on actin filaments but also on microtubules.

### Increased Auxin Contents in Intracellular Space Does not Inhibit PIN2 Endocytosis

Auxin has been suggested to repress the endocytosis of PINs-related carriers (Paciorek et al., 2005). Therefore, we firstly mimicked higher auxin contents in intracellular space in transgenic *DII-VENUS* materials via NPA action (Figure 2C). Our results showed that 10 μM NPA-induced weakened *DII-VENUS* signals in both B-supplied and B-deprived plants, and Al exposure further diminished it in the whole root tips (Figure 2C), suggesting it was feasible to enhance cellular auxin contents by NPA supply. Then we tested whether NPA-induced higher auxin in intracellular space inhibits PIN2 endocytosis. Our results shown that, in the absence of Al, NPA-induced increase of auxin contents in cellular spaces did not block PIN2 endocytosis in the presence of BFA (Figure S5). Moreover, BFA-induced PIN2 spots were also detected even in Al-treated plants applied with B and NPA. These results show that Al-induced increase of cellular auxin contents does not inhibit PIN2 endocytosis.

### B Supply Promotes PIN2 Carriers Turnover and Ameliorates Al Effects

To give further insight into B functions and Al effects on PIN2 endocytosis, we employed FM4-64 dye to label the plasma membrane of root apices of transgenic *PIN2-GFP* plants at various ratios of BFA, B and Al. Contrast distribution of the endocytic endosome labeled by FM4-64 and/or PIN2-GFP signals in the cytosol was found in Al-treated and Al-untreated roots (Figure 6A). In the absence of Al, BFA-induced spots labeled by both FM4-64 (red signals) and PIN2-GFP (green signals) were observed in the cytosol (Figure 6A), and the size of spots was bigger in B-supplied plants than that in B-deprived plants, suggesting the turnover of PIN2 with membrane vesicles which was promoted by B supply observed in Figure 3B. In the presence of Al, intensified FM4-64 signals in the endosomal membranes were observed in the cytosol, but the extent was higher in B-deprived plants than that in B-supplied plants, implying Al-intensified the endocytosis of plasma membranes which was further elevated by B deprivation. However, in the presence of Al, those endosomes in the cytosol only labeled by FM4-64 signals but not by PIN2-GFP signals in B-deprived plants, while B-supplied plants also had some small spots labeled by both FM4-64 and PIN2-GFP signals. Relatively less endosomes could be congregated into BFA-induced spots in the presence of Al especially at B deprivation. Those results disclose that the endocytosis of plasma membranes, but without loading of PIN2 proteins, operates exhaustively after Al exposure which was exacerbated by B deprivation. Collectively, these results suggest that B supply specifically promoted PIN2 endocytosis to counteract the negative effects of Al under B deprivation conditions.

**Figure 6.**
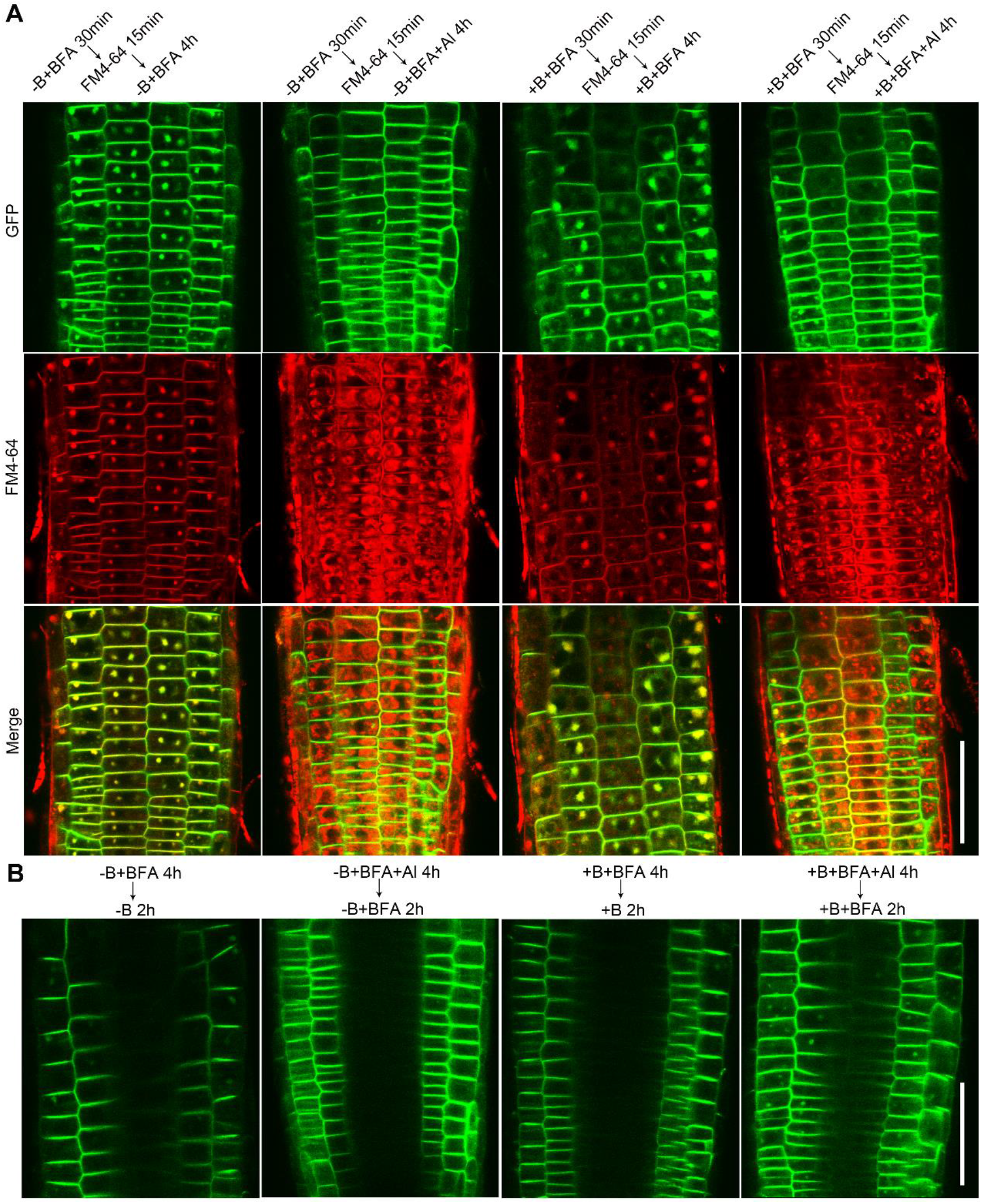
Effects on Al and B on the endocytosis and exocytosis of PIN2 carriers in the transition zone of *Arabidopsis* root tips. Six-day-old transgenic *PIN2-GFP* seedlings were treated with solution containing B [0 or 50 μM B(OH)_3_)] and BFA (0 or 100 μM) at pH 4.5 for 30 min, and further stained with 5 μM FM4-64 for 15 min. Plants were then exposed to solution containing Al (0 or 100 μM AlCl_3_), B [0 or 50 μM B(OH)_3_)] and BFA (0 or 100 μM) at pH 4.5 for 4 h. (A) Green and red colors indicate PIN2-GFP signals and FM4-64, respectively; merge images are the overlay of GFP and FM4-64 signals. **(B)** Effects of B and Al on the restoration and re-endocytosis of PIN2 carriers in *Arabidopsis* roots. Six-day-old *Arabidopsis* (*PIN2-GFP*) seedlings were treated with solution containing Al (0 or 100 μM AlCl_3_), B [0 or 50 μM B(OH)_3_)] and BFA (0 or 100 μM) at pH 4.5 for 4 h. Plants were then transferred to solution containing B [0 or 50 μM B(OH)_3_)] and BFA (0 or 100 μM) at pH 4.5 for 2 h. Thereafter, transition zones of treated roots (> 10) were imaged using CSLM. Bars are 100 μm of **(A)** and **(B)**.

In order to decipher the effects of B and Al on the exocytosis and recovery of the endocytosis of PIN2 carriers in the TZ, we compared the disappearance rate of BFA-induced PIN2 spots in both B-supplied plants and B-deprived plants. In the absence of Al, the disappearance rate of BFA-induced PIN2 spots was faster in B-sufficient plants than in B-deprived plants (Figure 6B). These results suggest that B supply promoted the exocytosis of BFA-induced PIN2 spots. For the analysis of recovery of the endocytosis of PIN2 proteins which were inhibited by Al, we compared the formation rate of BFA-induced PIN2 spots once Al action was interrupted. Again, we have detected some BFA-induced PIN2 spots in B-supplied but not in B-deprived plants after 2 h of Al removal. These results demonstrate that, in addition to endocytosis, B supply also promoted the exocytosis of PIN2 proteins.

### B Supply Promotes the Trafficking of PIN2-dependent IAA Transport to Reverse Al Inhibition

According to the typical chemiosmotic hypothesis, PIN2 membrane proteins execute their functions on auxin efflux from cells, but this mechanism cannot explain the Al-induced enhancement of PIN2 membrane signals in the root apical TZ along with decrease of auxin concentration in the EZ. A plausible model has been proposed that IAA is stored in endosomes by vesicular transporters and then secreted via endosomal exocytosis like a neurotransmitter secretory pathway (Schlicht et al., 2006). In order to validate this suggestion, we analyzed the spatial distribution of IAA marked by specific antibody in the transgenic *PIN2-GFP* seedlings. Our results revealed that, in the absence of Al, except some diffuse cytoplasmic signals, IAA was mainly located in the plasma membranes, especially in the horizontal plasma membranes of root apical TZ, regardless of B supply (Figure 7A and 7B). BFA exposure not only facilitated the formation of a large PIN2 spots in cytoplasmic spaces (Figure 7A), but also depleted IAA signals at the plasma membranes along with scored them into BFA-induced PIN2 spots, in both B-deprived and -sufficient plants. It indicates the transport of IAA adopts the vesicle trafficking of PIN2 carriers-based. As observed in the enlarged Figure 7B, invaginated membrane regions seemingly were coated by clathrin, which mediates the endocytosis of PIN2-based auxin transport. In the presence of Al, Al exposure significant intensified both IAA and PIN2 membrane signals (Figure 7A, 7C and 7D). Moreover, IAA signals were scored within BFA-induced PIN2 spots in B-supplied plants even in the presence of Al. Collectively, we suggest that Al-induced inhibition of polar auxin transport from the TZ to EZ via repressing the trafficking of PIN2 membrane carriers, while this adverse effect can be alleviated by B supply.

**Figure 7.**
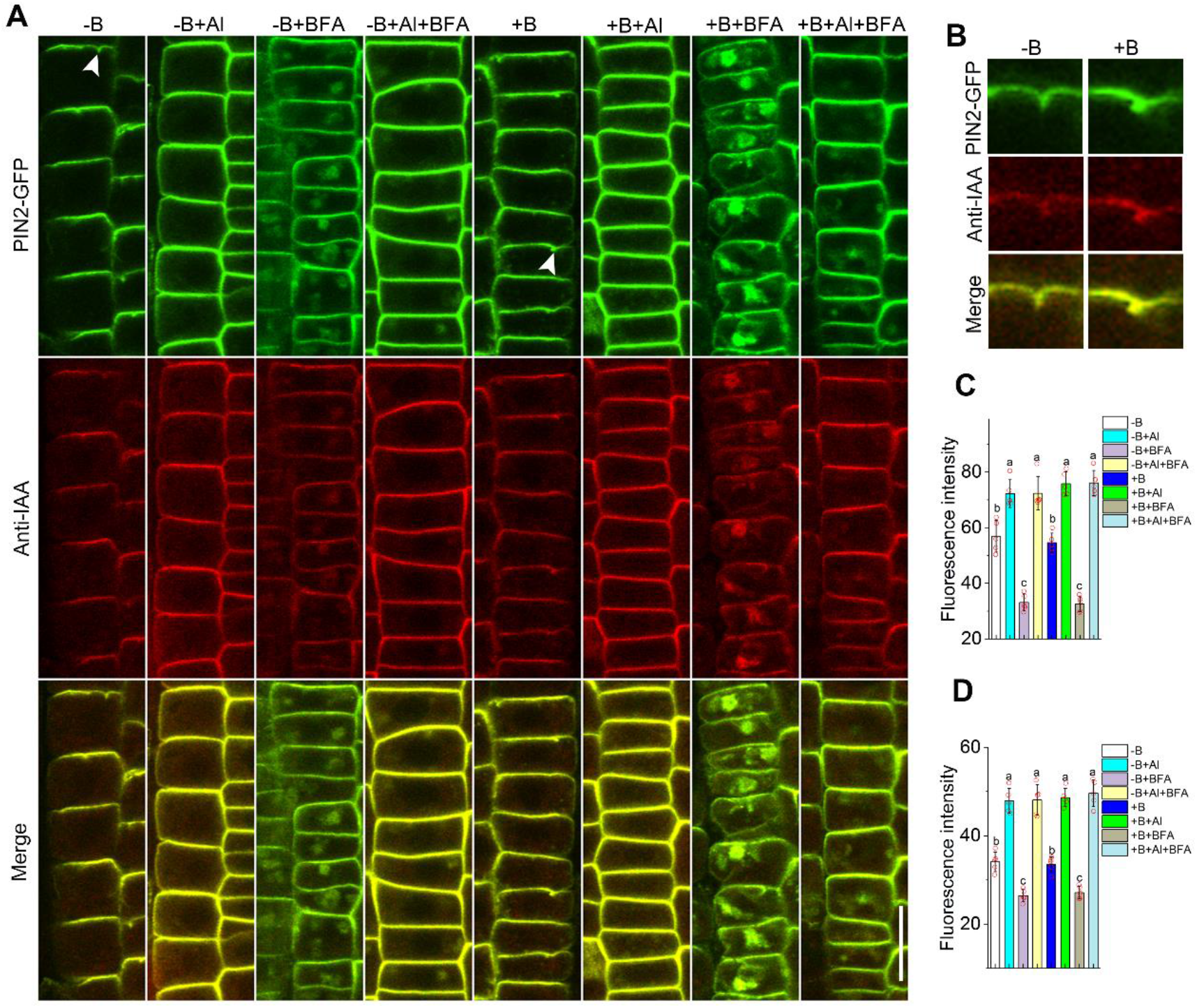
Colocalization of PIN2 carriers and auxin in cells of root apex transition zones of transgenic PIN2-GFP seedlings. Six-day-old transgenic *PIN2-GFP* seedlings were further treated with solution containing Al (0 or 100 μM AlCl_3_), B [0 or 50 μM B(OH)_3_)] and BFA (0 or 100 μM) at pH 4.5 for 4 h. **(A)** Green and red colors indicate PIN2-GFP signals and IAA signals, respectively; merge images are the overlay of GFP and IAA signals. **(B)** Images which indicated by white arrows in **(A)** are enlarged. **(C)** and **(D)** are the fluorescence intensity of PIN2-GFP and IAA signals at the plasm membrane, respectively. Values are means ± SD (*n* = 5). Red circles represent individual data. Data labelled with different letters are significantly different at P < 0.05 (LSD’s test). Bar is 20 μm in **(A)**.

### Al-Induced Inhibition of Root Growth Independently of Auxin Signaling

To link detrimental effects of Al on root growth with auxin signaling, we analyzed root growth in the presence of auxinole (Ax), a potent auxin antagonist of TRANSPORT INHIBITOR RESPONSE 1/AUXIN SIGNALLING F-BOX (TIR1/AFB) receptor that binds with TIR1 to block the formation of TIR1/AFB-IAA-AUX/IAA complex. As mentioned of transgenic *DII-VENUS* in Brunoud et al. (2012), fluorescent proteins of VENUS label with DII domain of AUX/IAA repressors. Because Ax can block the formation of TIR1/AFB-IAA-AUX/IAA complex, so fluorescent proteins of VENUS cannot be degraded. Before doing root growth test, we first analyzed transgenic *DII-VENUS* materials to verify 25 μM Ax could efficiently block auxin signaling (Figure 8A).

**Figure 8.**
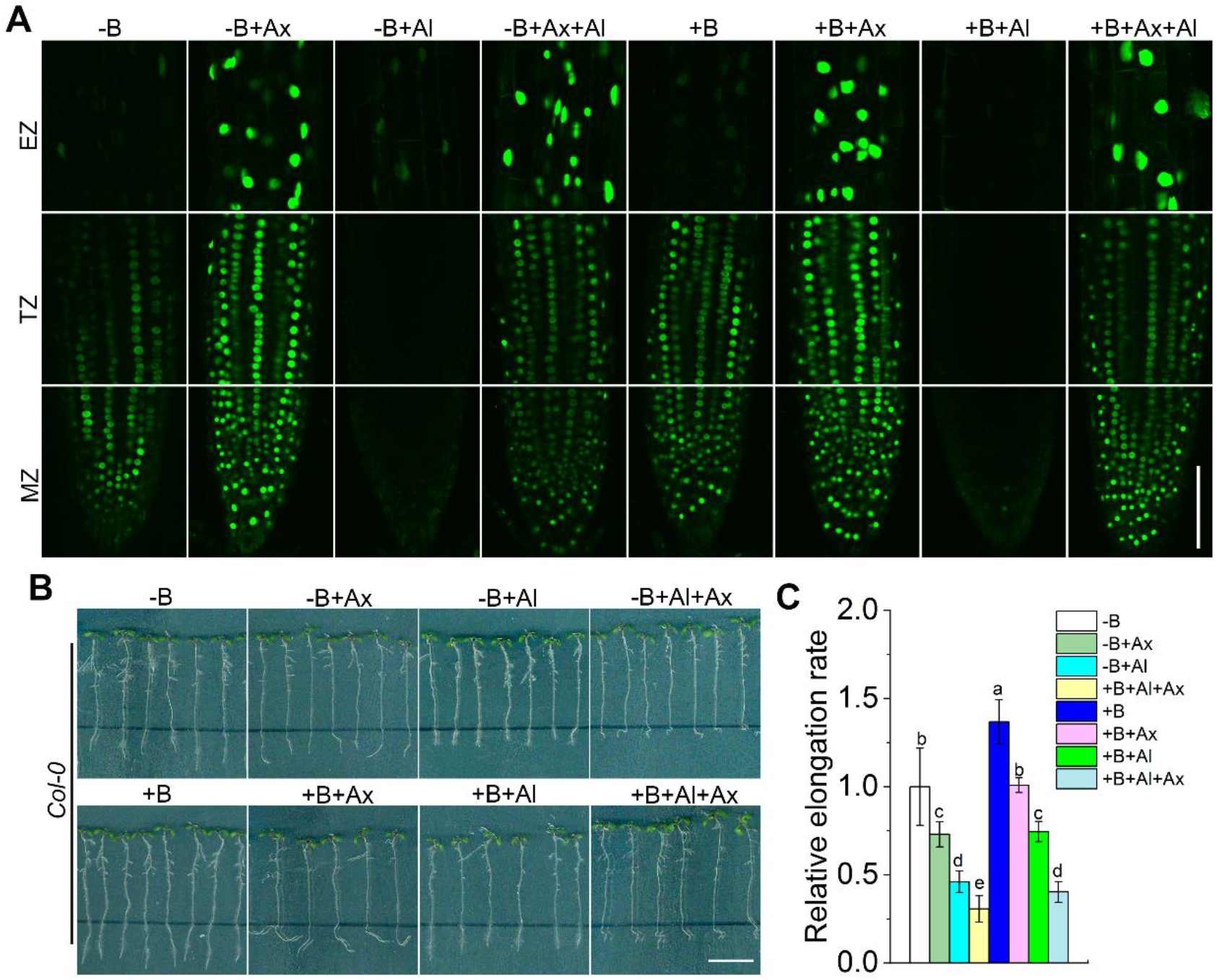
Disruption of auxin signaling by auxinole does not alleviate Al-induced inhibition of root growth. Six-day-old seedlings of transgenic *DII-VENUS* seedlings were treated with solution containing Al (0 or 100 μM AlCl_3_), B [0 or 50 μM B(OH)_3_)] and Ax (0 or 25 μM auxinole) at pH 4.5 for 4 h. **(A)** one (of > 10) typical CSLM images for each treatment.(A) Wild type (*col-0*) plants were grown on agar containing for 6 days that then exposed to Al (0 or 100 μM AlCl_3_), B [0 or 50 μM B(OH)_3_)] and Ax (0 or 25 μM auxinole) at pH 4.5 for 2 d. The black lines indicate the original locations of root tips at beginning of treatment. Subsequently, treated seedlings (> 20) were imaged and relative elongation rate **(C)** was calculated and -B treatment was set to 1. **(C)** Values are means ± SD (*n* = 10); red circles represent individual data. Bars are 100 μm for **(A)** and 1 cm for **(B)**, respectively.

As expected, Ax-induced increase of DII-VENUS signals in both B-deprived and - sufficient plants as well as the in the entire root tips with little difference among the root zones, but this elevation was lower in the MZ and TZ of Al-treated plants (Figure 8A). This implies that Al-induced increase in auxin content, in the intracellular space of the epidermal cells in root tips, possible facilitated AUX/IAA repressors marked by fluorescent VENUS proteins to be targeted into vacuoles for lytic degradation. Thereafter, we further assayed wild type (*col-0*) root growth in the presence of Ax. Our results show that Ax exposure not only blocked auxin signaling, but also repressed root gravitropic responses and growth elongation, regardless of B and/or Al supply (Figure 8B). Interestingly, Al-induced inhibition of root growth had not been alleviated via Ax application (Figure 8C). These results suggest that Al-induced inhibition of root growth depends on polar auxin transport pathway rather than on auxin signaling.

## Discussion

Boron has been reported to ameliorate Al-induced inhibition of root growth via differential processes such as decreasing Al accumulation in the cell wall pectin (Yu et al., 2009; Stass et al., 2007), alkalization of apoplast pH in root apical transition zone (Li et al., 2018) and strengthening antioxidant responses (Corrales et al., 2008). At the same time, either B deprivation or Al stress can repress root growth, via modulating polar auxin transport in root tips (Yang et al., 2014; Sun et al., 2010; Camacho-Cristóbal et al., 2015). This work has elucidated the mechanistic basis of this process and interaction between two elements. We have shown that B supply effectively diminishes auxin accumulation in the MZ caused by Al (Figure 2A and 2C) and traffic it to EZ instead (Figure 2A) that is essential for cellular growth and development (Teale et al., 2006).

Al-induced auxin accumulation in the root apical TZ is regarded as an important factor for root growth inhibition (Yang et al., 2014), but perturbation of auxin signaling pathways by auxin antagonist of Ax does not alleviate Al-induced inhibition of root growth (Figure 8B and 8C). Those results display that root growth inhibition by the increase of auxin content in root apices under Al stress is possibly via a non-transcriptional branch of TIR1/AFB signaling, as suggested by Fendrych et al. (2018). Similar to cold stress-induced inhibition of root growth that depends on repressing the endocytosis of PIN2-based auxin transport from the MZ to the EZ, but independent on auxin signaling (Shibasaki et al., 2009). It seems that the capacity to circulate auxin through PINs carriers regulates the root growth (Billou et al., 2005). Thus, we suggest that Al-induced inhibition of root growth depends on auxin transport rather than auxin signaling.

Auxin transport-related carriers play a critical role in deploying and transporting auxin during root development (Petrášek and Friml, 2009; Friml, 2010). Acropetal auxin transport is regulated by auxin efflux carriers such as PIN1, PIN3, PIN4 and PIN7 (Petrášek and Friml 2009). In this study, we have shown that Al-induced increase of PIN1-GFP signals depended on elevated *PIN1* gene expression in the MZ (Figure 3A and S4); this result may underpin shoot-derived auxin transport toward root apices. As observed in Figure 2C, NPA treatment efficiently diminished root apical auxin content even in the presence of Al. In contrast, the apically localized PIN2 in the epidermal cells is responsible for the basipetal auxin transport (Baluška et al., 2010). Al-intensified PIN2-GFP membrane signals in root apical TZ (Figure 3B; Shen et al., (2008)) indicates more PIN2 carriers locate in plasma membrane under Al exposure. The pharmacological experiments using CHX and COR to inhibit the transcription and protein biosynthesis of *PIN2* showed that Al-induced increase in PIN2 membrane signals does not depend on either transcription or protein biosynthesis of *PIN2*, but the inhibition of PIN2 endocytosis (Figure 4A). Moreover, the result of Al-caused decrease of auxin content in root apical EZ of B-deprived plants (Figure 2A) hints that PM-resident PIN2 carriers are less efficient in performing auxin efflux in the presence of Al. It is similar to the research on PIN1 carriers, which still locate at the plasma membrane after the treatment of BFA or auxin transport inhibitor, but displays the physiological effects of the inhibition of auxin efflux (Delbarre et al., 1998; Geldner et al., 2001; Rahman et al., 2007). It is plausible that Al-blocked endocytosis of PIN2 membrane carriers to repress the basipetal auxin transport from root apical MZ to EZ.

Auxin transporters undergo continuous endo- and exo-cytosis between the plasma membranes and endosomes (Adamowski and Friml, 2015). Moreover, it has been reported that metal ions stresses (Mn, Fe, Cu, Cd and Al), disrupted auxin biosynthesis and distribution in root apices. Intriguingly, differential metal ions interfered with different auxin transport-related carriers. Mn toxicity-induced decrease of auxin content in root apices was related to downregulating the expression of *PIN4* and *PIN7* genes (Zhao et al., 2017); Cu stress-mediated auxin redistribution was associated with modulating PIN1 carriers rather than PIN2 carriers (Yuan et al., 2013); Cd toxicity was causally related to reduce the abundance of PIN1/3/7 membrane proteins and not associated with transcriptional regulation (Yuan and Huang, 2016) and Fe stress-downregulated PIN2 expression to affect root growth (Li et al., 2015). In this study, we show that Al exposure resulted in a specific repression of the endocytosis of PIN2 membrane proteins (Figure 2B) and upregulation of PIN1 gene expression (Figure 3A and S4) in MZ. Altogether, these studies have demonstrated that auxin transport-related carriers display diverse sensitivity to differential metal ions stress, including Al toxicity. Our results have revealed that Al toxicity selectively targeted and blocked endocytosis of PINs-efflux carriers such as PIN2; while other auxin carriers, including PIN1, PIN3, PIN7 and AUX1 proteins at the plasma membrane remain unaffected (Figure 2A, 2B and S1A). Thus, we suggest Al stress specific represses the endocytosis of PIN2 membrane carriers in the TZ to interrupt auxin transport toward the root apical EZ.

It has been reported that Al interfered with FM4-64 internalization and repressed the formation of BFA-induced compartments (Illéš et al., 2006) and inhibited the endocytosis of PIN2 membrane proteins (Shen et al., 2008) in *Arabidopsis* root apex transition zone. As mentioned above, we have observed that B supply promoted the endocytic behavior such as the endocytosis of PIN2 membrane proteins (Figure 3B and 6A) to reverse Al inhibition. Moreover, Al-intensified the endocytosis of plasma membrane but without PIN2 carrier loading in B-deprived roots further suggests Al blocks the endocytosis of PIN2 membrane carries (Figure 6A). Despite the formation of clathrin-coated vesicles has been suggested to play critical role in mediating the internalization of plasma membrane-resident PIN proteins and their polar localization (Dhonukshe et al., 2007; Kitakura et al., 2011; Wang et al., 2013). Whether Al stress affects clathrin-coated process will need more evidences from future studies.

Actin filaments play critical role in many processes such as endocytosis, vesicle motility and auxin transport (Dhonukshe et al., 2008), and actin-based vesicle trafficking of PINs carriers has been suggested in previous reports (Shen *e*t al., 2008; Marhavý et al., 2011; Shibasaki et al., 2009). Interestingly, actin stabilization by auxin transport inhibitors such as 2,3,5-triiodobenzoic acid (TIBA) and 2-(1-pyrenoyl) benzoic acid (PBA) can repress PIN endocytosis along with disrupting of auxin efflux out of cells and auxin transport (Geldner et al., 2001; Rahman et al., 2007). As mentioned above, our results display that Al-induced actin stabilization (Figure 5A), repression of PIN2 endocytosis (Figure 4A) and blocking of polar auxin transport (Figure 2A), and it is plausible that Al stress executes its negative action via stabilizing actin-based PIN2 endocytosis to repress polar auxin transport (Figure 5E). Thus, the depolymerization of actin filaments by LatB could disrupt endocytic pathways of PIN2 carriers (Figure 5E). Similar to Al-caused changes of actin filaments, perturbation of microtubules by Al stress (Figure 5C) or the depolymerization of microtubules by Oryz (Figure 5E) also represses PIN2 endocytosis. Therefore, we suggest that not only actin filaments but also microtubules are required for PIN2 endocytosis.

According to conventional view, polar auxin transport (PAT) is executed by auxin transported-related carriers such as PINs when they reside at the plasma membranes. However, this model is failing to explain why auxin accumulates within vesicles and endosomes (Šamaj et al., 2004; Schlicht et al., 2006; Mettbach et al., 2017; Baluška et al., 2018; Pařízková et al., 2021). Moreover, this model also does not explain why 20 min of BFA exposure causes a rapid inhibition of the basipetal auxin transport while PIN1 carriers still locate at the plasma membrane (Delbarre et al., 1998). In order to explain this controversy, a neurotransmitter-like mode for IAA secretion out of cells by recycling vesicles have been suggested (Schlicht et al., 2006; Mettbach et al., 2017; Baluška et al., 2018). It has been suggested that IAA in the cellular spaces was internalized into endocytic PIN2 endosomes, and then IAA was secreted out of cells by the fusion of endosomal recycling vesicles containing PIN2 carriers and IAA with the plasma membranes (Schlicht et al., 2006; Mettbach et al., 2017; Baluška et al., 2018). Here, we have further confirmed the plausibility of this model and showed that BFA exposure not only clustered the PIN2 membranes proteins, but also depleted and transported the plasma membrane IAA into endosomal vesicles (Figure 7A). B supplementation facilitated both endocytosis and exocytosis of PIN2 carriers in TZ even in the presence of Al (Figure 6A and 6B). Overall, B supply may restore Al-induced depletion in auxin content in the EZ by facilitating the vesicle trafficking of PIN2-based auxin transport in root apical TZ.

A new model is proposed to summarize our results (Figure 9). According to this model, Al-induced upregulation of PIN1 gene expression in the MZ (Figure 9A) may underpin stronger acropetal transport of shoot-derived auxin. At the meantime, Al stress selectively represses the endocytosis and recycling of PIN2 to hamper basipetal auxin transport form the TZ to EZ. This results in higher auxin accumulation in root MZ and TZ and triggers auxin deficiency in EZ. B supply downregulates PIN1 protein levels as well as reduces Al-binding with root tips (Figure 1C; Yu et al., (2009)) to diminish the impact of Al on PIN2 endocytosis-dependent polar auxin transport (Figure 9B). Overall, this study improves our understanding of how Al stress causes auxin accumulation in the meristem zone and transition zone while auxin starvation in the elongation zone in root apices, and suggests new mechanism of boron supply in acid soils in alleviating Al toxicity.

**Figure 9.**
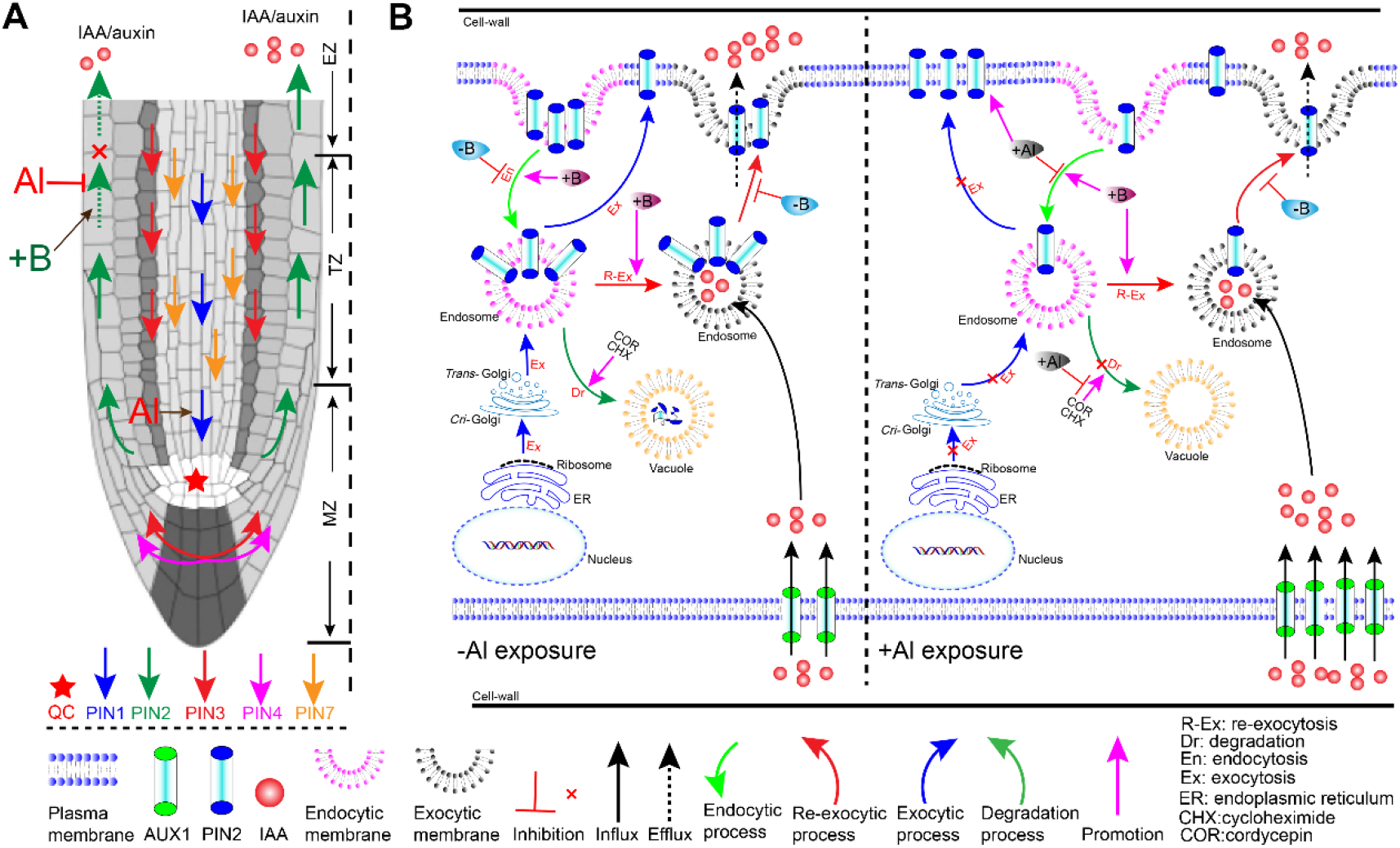
Suggested model depicting the routes of auxin transport in *Arabidopsis* root and their regulation by boron /Aluminum. **(A)** Auxin transport from shoot to the root apical quiescent center (QC) depends on the co-operation of PIN1, PIN3, PIN4 and PIN7 carriers, located in the pericycle and stele. The subsequent auxin redistribution from QC to the elongation zone mainly depends on PIN2 carriers that reside at the upper plasma membrane of the epidermal cells. This process is inhibited by Al stress, ultimately leading to auxin accumulation in the root apical MZ (meristem) and TZ (transition) zones, with less auxin being transported to the elongation zone (EZ). Boron presence ameliorates this process. **(B)** A suggested model of auxin efflux from a cell in the epidermal root apices. The endocytic PIN2 proteins are co-localized with IAA at the endosomes, and the recycling of endosomes (containing PIN2 proteins and IAA) promotes auxin efflux. Al blocks the endocytosis of PIN2 proteins at post-transcriptional level. B supply not only promotes endocytosis of PIN2 carrier, but also enhances its re-exocytosis, to restore Al-induced auxin depletion in the EZ.

## Materials and Methods

### Plant Materials and Growth Conditions

All lines are in the Columbia background of *Arabidopsis thaliana*. The transgenic of *ABD2-GFP* (actin-binding domain 2 of fimbrin gene fused with the green fluorescent protein gene, Wang et al., 2008), *MAP4-GFP* (microtubule-associated protein 4 gene fused with green fluorescent protein gene, Rosero et al., 2013), *DII-VENUS* (Brunoud et al., 2012), *DR5-GFP* (Billou et al., 2005), *PIN1-GFP* (Benková et al., 2003) and *PIN2-GFP* (Billou et al., 2005) seeds originated from our laboratories. All *Arabidopsis* seeds were sterilized by 10 % sodium hypochlorite for 3 min and treated with 75 % ethanol for 2 min followed by a thorough rinsing with sterilized water for three times. The sterilized seeds were sown on 0.7 % (w/v) agar-solidified nutrition medium containing 1/2 Murashige and Skoog (MS) solution with added B (0 or 50 μM H_3_BO_3_) and 1% sucrose at pH 5.8. After sowing, the plates were incubated for 2 d at 4 °C in the dark. Subsequently, all seedlings were grown for 6 d in a growth chamber with 16 h light/ 8 h dark at 25 °C. Six-day-old seedlings were further exposed with differential treatments.

### Root Growth Experiments

After 6-d growth, selected seedlings of uniform length were transferred to a 0.7% (w/v) agar-solidified media containing Al (0 or 100 μM AlCl_3_), B (0 or 50 μM B(OH)_3_) and Ax (0 or 25 μM auxinole, a potent auxin receptor antagonist.) at pH 4.5 for continued 2 d growth. The original locations of root apices were marked with black lines, and then all images were acquired by scanner (Epson Expression 11000XL). Subsequently, the newly grown lengths of roots were measured and calculated by the Image-J software and Excel 2009, respectively. Each replication has at least 30 plants and all experiments were repeated three times.

### Analysis of Al Distribution by Morin Staining

Six-day-old WT seedlings were exposed with 100 μM AlCl_3_ applied with 0 or 50 μM B(OH)_3_ for 0, 30 min, 2 h and 4 h, respectively. Roots were then stained with 0.01 % (w/v) morin dye (that binds to active Al) for 5 min and washed 3 min with deionized water. Al-morin signals in various functional root zones were detected by confocal laser scanning microscopy (CLSM; FV1000, Olympus, Tokyo, Japan) using 40-fold objective lens. Excitation and emission wavelength of morin were 488 nm and 520 nm, respectively.

### Pharmacological Treatments and Microscopic Analysis

Six-day-old WT and transgenic seedlings were treated with 0.2 mM CaCl_2_ solution containing combination of Al (0 or 100 μM AlCl_3_) and B (0 or 50 μM B(OH)_3_) (pH 4.5) for 4 h. Depending on the purpose of experiment, the following chemicals were added: NPA (0 or 10 μM, an auxin efflux inhibitor), BFA (0 or 100 μM, an inhibitor for exocytosis), CHX (0 or 50 mM cycloheximide, an inhibitor for protein biosynthesis), COR (0 or 100 μM cordycepin, transcriptional inhibitor), LatB (0 or 10 μM latrunculin B, a reagent to disrupt actin filaments), Oryz (0 or 20 μM oryzalin, a reagent to disrupt microtubules) and Ax (0 or 25 μM auxinole, an auxin receptor antagonist). Subsequently, treated seedlings were analyzed by CLSM at 40-fold objective lens. Excitation and emission wavelength were 488 nm and 510 nm, respectively.

### Immunofluorescence Labeling

Six-day-old *Arabidopsis PIN2-GFP* seedlings were further treated with 0.2 mM CaCl_2_ solution containing 100 μM AlCl_3_ and/or 50 μM H_3_BO_3_ (pH 4.5) for 4 h. Excised apical root segments (1 cm long) were fixed with a solution containing 4% paraformaldehyde, 50 mM PIPES, 5 mM MgSO4 and 5 mM EGTA at pH 6.9 for 30 min at room temperature. Subsequently, samples were washed repeatedly with a phosphate buffer solution (PBS; 0.14 M NaCl, 2.7 mM KCl, 10.1 mM Na_2_HPO_4_, 1.8 mM K_2_HPO_4_, pH 7.4) for 5 min, and then transferred to PBS containing 0.2% bovine serum albumin (BSA) for 30 min. Next, samples were incubated with the primary antibodies of auxin (IAA) polyclonal antibodies diluted 1:20 for 2 h at a room temperature. After washing with PBS for 5 min, samples were incubated with the second antibody IgG conjugated with Alexa 568 for 2 h at a room temperature. Finally, all samples were detected with CLSM at 40-fold objective lens. Excitation and emission wavelength were 488 nm and 510 nm for GFP, and 543 nm and 610 nm for Alexa 568, respectively. At least 10 roots were recorded.

### Endocytic and Exocytic Experiments with PIN2 Carriers

For endocytic experiments, 6 d transgenic *PIN2-GFP* seedlings were treated with 0.2 mM CaCl_2_ solution containing B [0 or 50 μM B(OH)_3_)] and BFA (0 or 100 μM) at pH 4.5 for 30 min, and further stained with 5 μM FM4-64 for 15 min. Then Al treatment (0 or 100 μM AlCl_3_) was administered for 4 h.

For exocytic experiments, we compared the disappearing rate of BFA-induced PIN2 spots under different conditions. 6 d *Arabidopsis* (*PIN2-GFP*) seedlings were treated with 0.2 mM CaCl_2_ solution containing Al (0 or 100 μM AlCl_3_), B [0 or 50 μM B(OH)_3_)] and BFA (0 or 100 μM) at pH 4.5 for 4 h, subsequently, transferred to 0.2 mM CaCl_2_ solution containing B [0 or 50 μM B(OH)_3_)] and BFA (0 or 100 μM) at pH 4.5 for 2 h.

Fluorescent signals were detected from the transition zones of root tips by confocal laser scanning microscope (CLSM; FV1000, Olympus, Tokyo, Japan). Excitation and emission wavelength were 488 nm and 510 nm for GFP, and 543 nm and 610 nm for FM4-64, respectively.

### Statistical analysis

All experiments were repeated at least three times. All data were analyzed by SPSS (Statistic Package for Social Science) software and Excel 2009, and charts were made by Origin 8.0 software. Significance of difference was determined at LSD’s test (*P* < 0.05).

## Acknowledgments

The authors like to thank the anonymous reviewers for their valuable comments to improve the quality of this work.

## References

Adamowski M, Friml J (2015) PIN-dependent auxin transport: action, regulation, and evolution. Plant Cell 27: 20–32

Baluška F, Mancuso S (2013) Root apex transition zone as oscillatory zone. Front Plant Sci 4: 1–15

Baluška F, Mancuso S, Volkmann D, Barlow PW (2010) Root apex transition zone: a signalling-response nexus in the root. Trends Plant Sci 15: 402–408

Baluška F, Strnad M, Mancuso S (2018) Substantial evidence for auxin secretory vesicles. Plant Physiol 176: 2586–2587

Benková E, Michniewicz M, Sauer M, Teichmann T, Seifertová D, Jürgens G, Friml J (2003) Local, efflux-dependent auxin gradients as a common module for plant organ formation. Cell 115: 591–602

Billou I, Xu J, Wildwater M, Willemsen V, Paponov I, Frimi J, Heldstra R, Aida M, Palme K, Scheres B (2005) The PIN auxin efflux facilitator network controls growth and patterning in Arabidopsis roots. Nature 433: 39–44

Brunoud G, Wells DM, Oliva M, Larrieu A, Mirabet V, Burrow AH, Beeckman T, Kepinski S, Traas J, Bennett MJ, et al (2012) A novel sensor to map auxin response and distribution at high spatio-temporal resolution. Nature 482: 103–106

Camacho-cristóbal JJ, Martín-rejano EM, Herrera-rodríguez MB, Navarro-gochicoa MT, Rexach J, González-fontes A, Fisiología D De, Celular B, Olavide UP De (2015) Boron deficiency inhibits root cell elongation via an ethylene / auxin / ROS-dependent pathway in Arabidopsis seedlings. 66: 3831–3840

Corrales I, Poschenrieder C, Barceló J (2008) Boron-induced amelioration of aluminium toxicity in a monocot and a dicot species. J Plant Physiol 165: 504–513

Delbarre A, Muller P, Guern J (1998) Short-lived and phosphorylated proteins contribute to carrier-mediated efflux, but not to influx, of auxin in suspension-cultured tobacco cells. Plant Physiol 116: 833–844

Dhonukshe P, Aniento F, Hwang I, Robinson DG, Mravec J, Stierhof YD, Friml J (2007) Clathrin-mediated constitutive endocytosis of PIN auxin efflux carriers in Arabidopsis. Curr Biol 17: 520–527

Dhonukshe P, Grigoriev I, Fischer R, Tominaga M, Robinson DG, Hašek J, Paciorek T, Petrášek J, Seifertová D, Tejos R, et al (2008) Auxin transport inhibitors impair vesicle motility and actin cytoskeleton dynamics in diverse eukaryotes. Proc Natl Acad Sci U S A 105: 4489–4494

Eticha D, Stass A, Horst WJ (2005) Cell-wall pectin and its degree of methylation in the maize root-apex: significance for genotypic differences in aluminium resistance. Plant, Cell Environ 28: 1410–1420

Fendrych M, Akhmanova M, Merrin J, Glanc M, Hagihara S, Takahashi K, Uchida N, Torii KU, Friml J (2018) Rapid and reversible root growth inhibition by TIR1 auxin signalling. Nat Plants 4: 453–459

Friml J (2003) Auxin transport-shaping the plant. Curr Opin Plant Biol 6: 7–12

Friml J (2010) Subcellular trafficking of PIN auxin efflux carriers in auxin transport. Eur J Cell Biol 89: 231–235

Gasman S, Kalaidzidis Y, Zerial M (2003) RhoD regulates endosome dynamics through Diaphanous-related Formin and Src tyrosine kinase. Nat Cell Biol 5: 195–204

Geisler M, Murphy AS (2006) The ABC of auxin transport: The role of p-glycoproteins in plant development. FEBS Lett 580: 1094–1102

Geldner N, Friml J, Stierhof YD, Jürgens G, Palme K (2001) Auxin transport inhibitors block PIN1 cycling and vesicle trafficking. Nature 413: 425–428

Illéš P, Schlicht M, Pavlovkin J, Lichtscheidl I, Baluška F, Ovečka M (2006) Aluminium toxicity in plants: internalization of aluminium into cells of the transition zone in Arabidopsis root apices related to changes in plasma membrane potential, endosomal behaviour, and nitric oxide production. J Exp Bot 57: 4201–4213

Kitakura S, Vanneste S, Robert S, Löfke C, Teichmann T, Tanaka H, Friml J (2011) Clathrin mediates endocytosis and polar distribution of PIN auxin transporters in Arabidopsis. Plant Cell 23: 1920–31

Kollmeier M, Felle HH, Horst WJ (2000) Genotypical differences in aluminum resistance of maize are expressed in the distal part of the transition zone. Is reduced basipetal auxin flow involved in inhibition of root elongation by aluminum? Plant Physiol 122: 945–956

Kopittke PM, Moore KL, Lombi E, Gianoncelli A, Ferguson BJ, Blamey FPC, Menzies NW, Nicholson TM, McKenna BA, Wang P, et al (2015) Identification of the primary lesion of toxic aluminum in plant roots. Plant Physiol 167: 1402–1411

Li G, Song H, Li B, Kronzucker HJ, Shi W (2015) Auxin resistant1 and PIN-FORMED2 protect lateral root formation in arabidopsis under iron stress. Plant Physiol 169: 2608–2623

Li X, Li Y, Mai J, Tao L, Qu M, Liu J, Shen R, Xu G, Feng Y, Xiao H, et al (2018) Boron alleviates aluminum toxicity by promoting root alkalization in transition zone via polar auxin transport. Plant Physiol 177: 1254–1266

Marhavý P, Bielach A, Abas L, Abuzeineh A, Duclercq J, Tanaka H, Pařezová M, Petrášek J, Friml J, Kleine-Vehn J, et al (2011) Cytokinin modulates endocytic trafficking of PIN1 auxin efflux carrier to control plant organogenesis. Dev Cell 21: 796–804

Mettbach U, Strnad M, Mancuso S, Baluška F (2017) Immunogold-EM analysis reveal brefeldin a-sensitive clusters of auxin in Arabidopsis root apex cells. Commun Integr Biol 10: e1327105

O’Neill MA, Ishii T, Albersheim P, Darvill AG (2004) RHAMNOGALACTURONAN II: structure and function of a borate cross-linked cell wall pectic polysaccharide. Annu Rev Plant Biol 55: 109–139

Paciorek T, Zažímalová E, Ruthardt N, Petrášek J, Stierhof YD, Kleine-Vehn J, Morris DA, Emans N, Jürgens G, Geldner N, et al (2005) Auxin inhibits endocytosis and promotes its own efflux from cells. Nature 435: 1251–1256

Pařízková B, Žukauskaitė A, Vain T, Grones P, Raggi S, Kubeš MF, Kieffer M, Doyle SM, Strnad M, Kepinski S, et al (2021) New fluorescent auxin probes visualise tissue-specific and subcellular distributions of auxin in Arabidopsis. New Phytol 230: 535–549

Péret B, Swarup K, Ferguson A, Seth M, Yang Y, Dhondt S, James N, Casimiro I, Perry P, Syed A, et al (2012) AUX/LAX genes encode a family of auxin influx transporters that perform distinct functions during arabidopsis development. Plant Cell 24: 2874–2885

Petrášek J, Friml J (2009) Auxin transport routes in plant development. Development 136: 2675–2688

Rahman A, Bannigan A, Sulaman W, Pechter P, Blancaflor EB, Baskin TI (2007) Auxin, actin and growth of the Arabidopsis thaliana primary root. Plant J 50: 363–376

Rosero A, Žárský V, Cvrčková F (2013) AtFH1 formin mutation affects actin filament and microtubule dynamics in Arabidopsis thaliana. J Exp Bot 64: 585–597

Sabatini S, Beis D, Wolkenfelt H, Murfett J, Guilfoyle T, Malamy J, Benfey P, Leyser O, Bechtold N, Weisbeek P, et al (1999) An auxin-dependent distal organizer of pattern and polarity in the Arabidopsis root. Cell 99: 463–472

Šamaj J, Baluška F, Voigt B, Schlicht M, Volkmann D, Menzel D (2004) Endocytosis, actin cytoskeleton, and signaling. Plant Physiol 135: 1150–1161

Schlicht M, Strnad M, Scanlon MJ, Mancuso S, Hochholdinger F, Palme K, Volkmann D, Menzel D, Baluska F (2006) Auxin immunolocalization implicates vesicular neurotransmitter-like mode of polar auxin transport in root apices. Plant Signal Behav 1: 122–133

Schwechheimer C, Yalovsky S, Zársky V (2021) Auxin does not inhibit endocytosis of PIN1 and PIN2 auxin efflux carriers. Plant Physiol 186: 808–811

Shen H, Hou N, Schlicht M, Wan Y, Mancuso S, Baluska F (2008) Aluminium toxicity targets PIN2 in Arabidopsis root apices: effects on PIN2 endocytosis, vesicular recycling, and polar auxin transport. Chinese Sci Bull 53: 2480–2487

Shibasaki K, Uemura M, Tsurumi S, Rahman A (2009) Auxin response in Arabidopsis under cold stress: underlying molecular mechanisms. Plant Cell 21: 3823–3838

Shorrocks VM, Bureau M (1997) The occurrence and correction of boron deficiency. Plant Soil 193: 121–148

Sivaguru M, Liu J, Kochian L V. (2013) Targeted expression of SbMATE in the root distal transition zone is responsible for sorghum aluminum resistance. Plant J 76: 297–307

Stass A, Kotur Z, Horst WJ (2007) Effect of boron on the expression of aluminium toxicity in Phaseolus vulgaris. Physiol Plant 131: 283–290

Sun P, Tian QY, Chen J, Zhang WH (2010) Aluminium-induced inhibition of root elongation in Arabidopsis is mediated by ethylene and auxin. J Exp Bot 61: 347–356

Tanaka M, Fujiwara T (2008) Physiological roles and transport mechanisms of boron: perspectives from plants. Pflugers Arch Eur J Physiol 456: 671–677

Teale WD, Paponov IA, Palme K (2006) Auxin in action: signalling, transport and the control of plant growth and development. Nat Rev Mol Cell Biol 7: 847–859

Verbelen JP, De Cnodder T, Le J, Vissenberg K, Baluška F (2006) The root apex of Arabidopsis thaliana consists of four distinct zones of growth activities: meristematic zone, transition zone, fast elongation zone and growth terminating zone. Plant Signal Behav 1: 296–304

von Uexküll HR, Mutert E (1995) Global extent, development and economic impact of acid soils. Plant Soil 171: 1–15

Wang C, Yan X, Chen Q, Jiang N, Fu W, Ma B, Liu J, Li C, Bednarek SY, Pan J (2013) Clathrin light chains regulate clathrin-mediated trafficking, auxin signaling, and development in Arabidopsis. Plant Cell 25: 499–516

Wang YS, Yoo CM, Blancaflor EB (2008) Improved imaging of actin filaments in transgenic Arabidopsis plants expressing a green fluorescent protein fusion to the C- and N-termini of the fimbrin actin-binding domain 2. New Phytol 177: 525–536

Wu D, Shen H, Yokawa K, Baluška F (2014) Alleviation of aluminium-induced cell rigidity by overexpression of OsPIN2 in rice roots. J Exp Bot 65: 5305–5315

Yang Y, Hammes UZ, Taylor CG, Schachtman DP, Nielsen E (2006) High-affinity auxin transport by the AUX1 influx carrier protein. Curr Biol 16: 1123–1127

Yang Z-B, Geng X, He C, Zhang F, Wang R, Horst WJ, Ding Z (2014) TAA1-regulated local auxin biosynthesis in the root-apex transition zone mediates the aluminum-induced inhibition of root growth in Arabidopsis. Plant Cell 26: 2889–2904

Yu M, Shen R, Xiao H, Xu M, Wang H, Wang H, Zeng Q, Bian J (2009) Boron alleviates aluminum toxicity in pea (Pisum sativum). Plant Soil 314: 87–98

Yu Q, Hlavacka A, Matoh T, Volkmann D, Menzel D, Goldbach HE, et al (2002) Short-term boron deprivation inhibits endocytosis of cell wall pectins in meristematic cells of maize and wheat root apices. Plant Physiol 130: 415–421

Yuan H, Huang X (2016) Inhibition of root meristem growth by cadmium involves nitric oxide-mediated repression of auxin accumulation and signalling in Arabidopsis. Plant Cell Environ 1: 120–135

Yuan HM, Xu HH, Liu WC, Lu YT (2013) Copper regulates primary root elongation through PIN1-mediated auxin redistribution. Plant Cell Physiol 54: 766–778

Zhao J, Wang W, Zhou H, Wang R, Zhang P, Wang H, Pan X, Xu J (2017) Manganese toxicity inhibited root growth by disrupting auxin biosynthesis and transport in Arabidopsis. Front Plant Sci 8: 1–8

Zhao Y (2010) Auxin biosynthesis and its role in plant development. Annu Rev Plant Biol 61: 49–64

